# Reinforcing neuron extraction and spike inference in calcium imaging using deep self-supervised learning

**DOI:** 10.1101/2020.11.16.383984

**Authors:** Xinyang Li, Guoxun Zhang, Jiamin Wu, Yuanlong Zhang, Zhifeng Zhao, Xing Lin, Hui Qiao, Hao Xie, Haoqian Wang, Lu Fang, Qionghai Dai

## Abstract

Calcium imaging is inherently susceptible to detection noise especially when imaging with high frame rate or under low excitation dosage. We developed DeepCAD, a self-supervised learning method for spatiotemporal enhancement of calcium imaging without requiring any high signal-to-noise ratio (SNR) observations. Using this method, detection noise can be effectively suppressed and the imaging SNR can be improved more than tenfold, which massively improves the accuracy of neuron extraction and spike inference and facilitate the functional analysis of neural circuits.

---

Calcium imaging enables parallel recordings of large neuronal ensembles in living animals^1–4^ and offers a new possibility for deciphering information propagation, integration, and computation in neural circuits^5^. To obtain accurate neuron extraction and spike inference for downstream neuroscience analysis, high-SNR calcium imaging is desired. However, due to the paucity of fluorescence photons caused by low peak accumulations and fast dynamics of *in vivo* calcium transients^6,7^, calcium imaging is easy to be contaminated by detection noise (*i.e*. photon shot noise and electronic noise), especially in functional imaging where high temporal resolution is particularly important for analyzing neural activities^8^.

To capture sufficient fluorescence photons for high-SNR calcium imaging, the most direct way is to use high excitation dosage, but concurrent photobleaching, phototoxicity^9,10^, and tissue heating^11^ are detrimental for sample health and photosensitive biological processes, which limits the maximal excitation power for long-term *in vivo* imaging^12^. More effective strategies include using brighter calcium indicators^7,13^ and more sensitive photoelectric detectors^14^, but their performances are still largely restricted in photon-limited conditions such as dendritic imaging and deep-tissue imaging. Apart from these physical or biological approaches, data-driven methods are promising to offer an alternative solution to recover faithful signals from degraded recordings and reduce the photon budget of calcium imaging. As an intelligent signal processing technique, deep learning has been adopted by microscopists and achieved impressive performance in fluorescence imaging^15–18^. However, calcium transients are highly dynamic, non-repetitive activities and a firing pattern cannot be captured twice. Previous schemes for obtaining ground-truth images (*i.e*. clean images without noise contamination or high-SNR images with the same underlying scene) by extending integration time or averaging multiple noisy frames are no longer feasible, posing an entrenched obstacle for conventional supervised learning methods.

In this paper, we present DeepCAD, a self-supervised learning method for calcium imaging denoising by over tenfold SNR improvement without requiring any high-SNR observations for training. DeepCAD is based on the insight that a deep neural network can converge to a mean estimator even the target image used for training is another corrupted sampling of the same scene^19^. When looking at calcium imaging data, we explored the temporal redundancy of pervasive video-rate imaging and found that any two consecutive frames can be regarded as two independent samplings of the same underlying firing pattern, which can be used for training of denoising models. Furthermore, the input and output data are designed to be 3D volumes rather than 2D frames to fully exploit spatiotemporal information in the time-lapse stack. We show that such a 3D self-supervised method is extremely effective for calcium imaging denoising and even the subtlest calcium fluctuations induced by a single action potential (AP) can be restored from severely corrupted images. Finally, a Fiji-based plugin along with a pre-trained model were released to make our method easy to access and convenient to use.

The general principle of DeepCAD is schematized in Fig. 1a. For network architecture, we employed 3D U-Net^20^ to aggregate spatiotemporal information in multiple frames using 3D convolutional layers (Supplementary Fig. 1, Methods), which endows DeepCAD with better denoising capability than 2D architecture or classical methods (Supplementary Fig. 2). Benefiting from the self-supervised strategy, a single low-SNR stack of ~3500 frames is sufficient to be a complete training set. To generate the training set, two sub-stacks consisting of interlaced frames were split from the original low-SNR stack and 3D tiles were extracted from these sub-stacks for training (Supplementary Fig. 3). They contain approximate identical calcium transients when the original stack was imaged at near video rate, which is common for commercial or customized microscopes. After proper training, interpretable features can be learned (Supplementary Fig. 4) and the model can be applied to subsequent acquisitions without extra training (Fig. 1b). Although the network was trained on specified spatial and temporal resolution, we found that it had non-inferior performance on various frame rates (Supplementary Fig. 5) and magnifications (Supplementary Fig. 6), indicating the great scalability and generalization for versatile applications of DeepCAD.

**Fig. 1.**
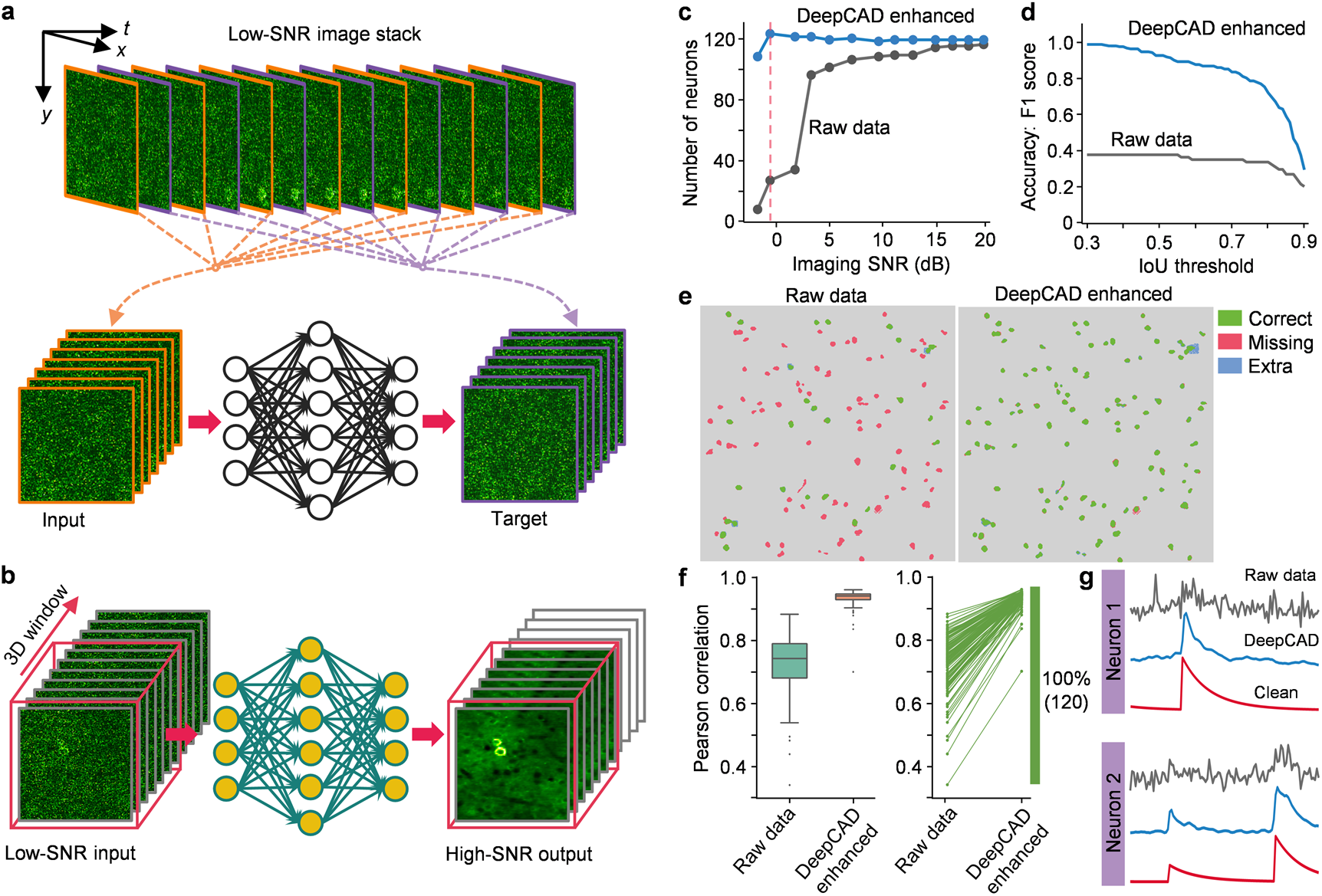
General principle and validation of DeepCAD. **a**, Self-supervised training strategy of DeepCAD. Consecutive frames in the original low-SNR stack are divided into two sub-stacks, used as the input volume and corresponding target volume to train a deep neural network (3D U-Net). After training, a denoising model can be established and memorized in network parameters. **b**, Application of the DeepCAD model. For subsequent acquisitions, a 3D (*x-y-t*) window traverses the entire stack and 3D tiles are sequentially fed into the pre-trained model. Denoised recordings will be obtained after the processing of the model. **c**, The number of neurons extracted under different imaging SNRs before and after the enhancement of DeepCAD. N=120 active neurons were simulated in the field of view (FOV). **d**, Accuracy of neuron segmentation quantified with F1 score at different intersection-over-union (IoU) thresholds (imaging SNR=-0.7 dB, indicated by the red dashed line in **c**). **e**, Spatial profiles of extracted neurons (imaging SNR=-0.7 dB). Correctly segmented regions (true positive) are colored green. Missing (false negative) and extra regions (false positive) are colored red and blue, respectively. Neuron extraction was implemented with CNMF^21^. **f**, Left: boxplot showing the distribution of Pearson correlation coefficients with clean traces before and after denoising (N=120). Right: increases of trace correlations. Each line represents one of 120 calcium traces and correlation coefficients of all neurons were observed improved. **g**, Calcium transients indiscernible from noise (gray) can be restored by DeepCAD (blue). Traces without noise contamination (red) serve as the ground truth for comparison.

To quantitatively evaluate the performance of DeepCAD, we first validated it on simulated calcium imaging data of different imaging SNRs (Supplementary Figs. 7-8 and Supplementary Notes 1-2), which contains synchronous noise-free recordings as the ground truth for comparison. The constrained nonnegative matrix factorization (CNMF) algorithm^21^ was used for downstream neuron extractions (Methods). After the enhancement of DeepCAD, more active neurons can be detected, especially when imaging SNR is low (Fig. 1c). The accuracy of neuron extraction was also quantified with F1 score and significant improvement was observed across a wide range of intersection-over-union (IoU) thresholds (Fig. 1d,e). For a typical IoU threshold of 0.7, the segmentation accuracy was improved by 2.4 folds (0.84 contrast to 0.35). Benefiting from the improved imaging quality, calcium traces extracted from the denoised data possess higher fidelity. To investigate the temporal enhancement of DeepCAD, we extracted calcium traces of all neurons from both raw noisy data and the enhanced counterpart. The Pearson correlation with the clean traces was significantly improved after denoising (Fig. 1f). Even the slightest calcium transients can be restored from the original noisy data (Fig. 1g and Supplementary Fig. 9). These facts suggest that the spatiotemporal enhancement of DeepCAD can improve the accuracy of neuronal localization and trace extraction and largely facilitate the analysis of neural circuits.

To verify the effectiveness and reliability of DeepCAD on neuroscience research, we then demonstrated its performance on two-photon calcium imaging based on data released by Svoboda lab^7^. In this dataset, simultaneous cell-attached electrophysiological recordings (Fig. 2a) are synchronized with two-photon imaging and serve as the reference of calcium transients and the ground truth of spike inference. Contaminated by detection noise, both the spatial footprint and temporal traces of the neuron were severely corrupted in the original data (Fig. 2b). After we applied DeepCAD to enhance these data, the annular cytoplasm became recognizable and calcium traces were liberated from noise (Fig. 2c and Supplementary Video 1). Even the most imperceptible calcium transients evoked by one AP, two APs, and three APs were clearly distinguished and still maintain their original dynamics (Fig. 2d-g), which otherwise would be submerged in noise. For further comparison, we extracted single-pixel fluorescence from cytoplasmic pixels and found that calcium transients can be unveiled at a single-pixel scale (Supplementary Fig. 10). Moreover, we performed spike inference (Methods) on traces extracted from the original data as well as the corresponding denoised data. Owing to the improvement of imaging SNR, the error rate of spike inference was consequently decreased (Fig. 2h and Supplementary Fig. 11). Among 107 independent calcium traces, 86% of them were observed to have lower error rates.

**Fig. 2.**
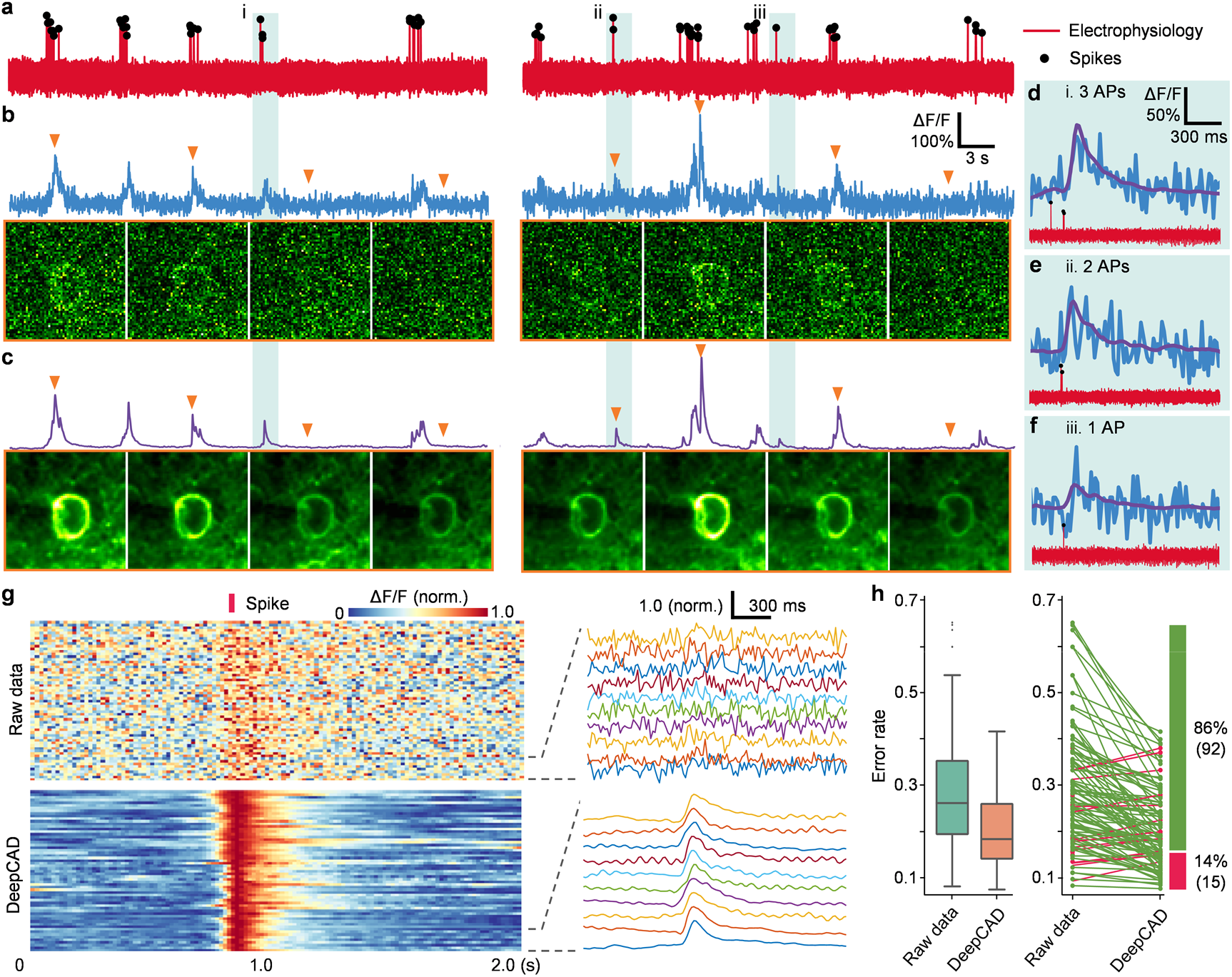
Spatiotemporal enhancement of DeepCAD. **a**, Electrophysiological recordings of neural activities from a single neuron. Detected spikes are marked with black dots. **b**, Two-photon calcium imaging of the same neuron synchronized with cell-attached electrophysiology. Both spatial footprints and temporal traces of the neuron were severely corrupted in detection noise. Representative frames indicated with orange triangles are presented below the trace. **c**, Fluorescence traces and representative frames after the enhancement of DeepCAD. **d-f**, The most imperceptible calcium transients evoked by three APs (**d**), two APs (**e**), and one AP (**f**) can be resolved and still keep their original dynamics noise removal. **g**, Calcium fluctuations evoked by 61 isolated action potentials. All spikes were normalized and temporally aligned with the red bar. Zoom-in traces are shown in the right panel. **h**, Left: Boxplot showing the distribution of error rates (lower is better) of spike inference for calcium traces extracted from enhanced data compared with those extracted from the original data (N=107). Real spike timings were revealed by simultaneous cell-attached recordings. Right: decreases of the error rate of spike inference. Each line represents one of 107 recordings, using green for decreased error rates and red for increased error rates.

Next, we employed DeepCAD for noise removal of calcium imaging of large neuronal populations in awake mice. To obtain high-SNR recordings for validation of our method, we designed and built a two-photon imaging system with the capability of simultaneous low-SNR and high-SNR recording (Supplementary Fig. 12 and Methods). The high-SNR detection path was strictly synchronized with the low-SNR detection path but with about 10-fold higher imaging SNR (Supplementary Fig. 13), which can be used as the reference for our denoising results. We first imaged spontaneous neuropil activities in cortical layer 1 of a transgenic mouse expressing GCaMP6f and found that calcium fluctuations indiscernible in original low-SNR recordings can be effectively recovered by DeepCAD (Fig. 3a-c and Supplementary Video 2). The imaging SNR was improved more than 10 folds considering that the SNR of enhanced recordings even surpasses corresponding high-SNR reference. Fluorescence traces of dendritic pixels can be accurately resolved and keep high consistency with the high-SNR reference (Fig. 3d-e and Supplementary Fig. 14). We also applied DeepCAD to enhance calcium imaging of somatic signals. After denoising, neuronal distribution and circuit dynamics can be recognized from a single frame (Fig. 3f-h and Supplementary Video 3). Using CNMF as the downstream source extraction method, 52.6% (229 contrast to 150) more active neurons can be extracted (Fig. 3i,j and Supplementary Fig. 15) and the trace peak SNR of extracted neurons was also improved more than two folds (9.9 contrast to 4.8, median value) (Fig. 3k), indicating that the functional analysis of large neuronal populations can be effectively strengthened due to improved SNR.

**Fig. 3.**
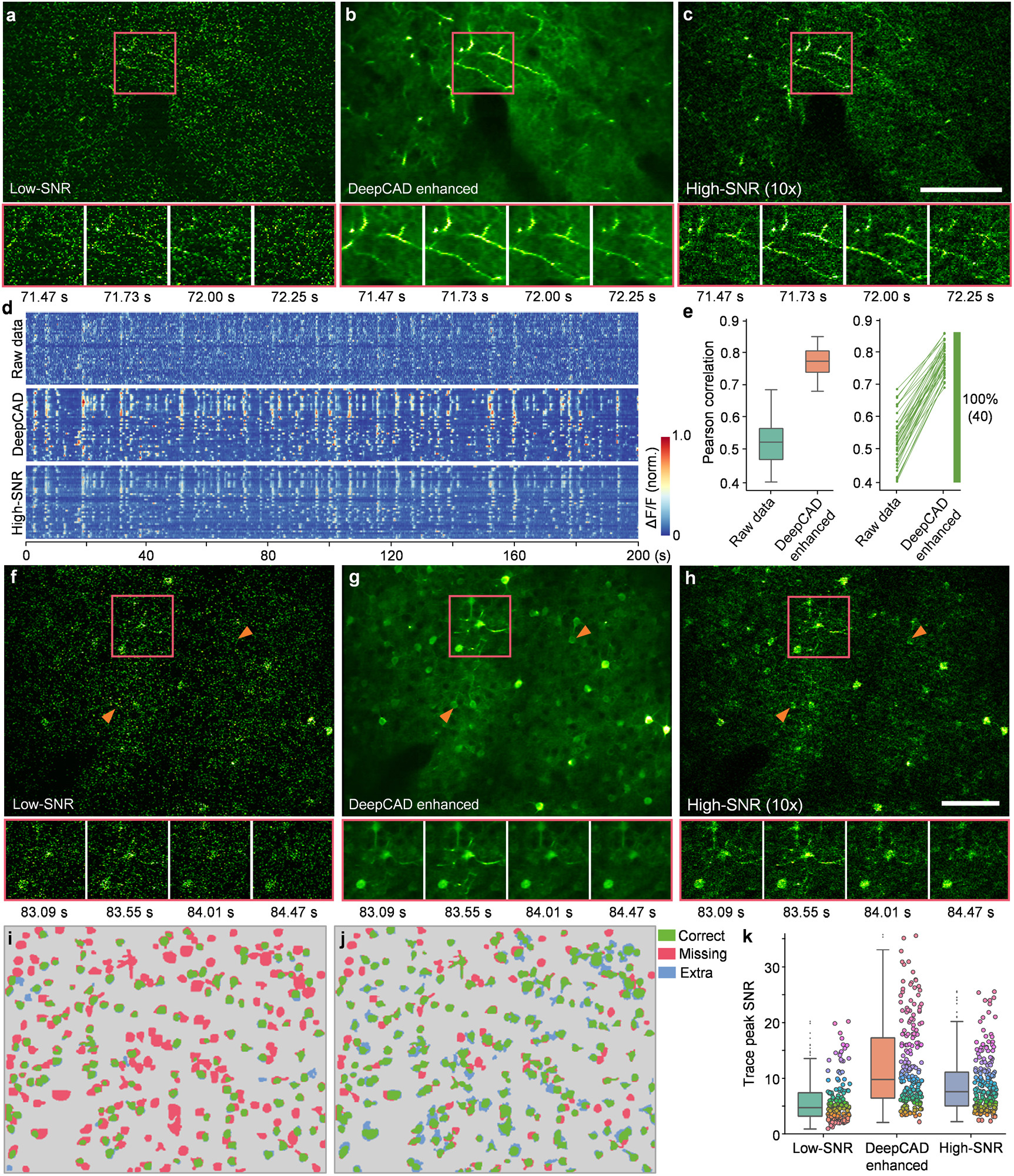
DeepCAD reinforces the recording of large neuronal populations. **a**, Spontaneous neuropil activities in layer 1 of the mouse cortex captured by the low-SNR detection path. **b**, Images restored from the low-SNR recording using DeepCAD. **c**, Synchronized recording acquired by the high-SNR detection path (10-fold imaging SNR). Magnified views of the boxed regions show calcium transients in a ~0.8 s time window. Scale bar 100 μm. **d**, Fluorescence traces extracted from 40 dendritic pixels. Top: low-SNR recording, Middle: DeepCAD enhanced recording, Bottom: high-SNR recording. **e**, Pearson correlation coefficients of single-pixel calcium traces before and after denoising (left). High-SNR traces were used as the reference for correlation calculation. Improvements were observed in all 40 traces (right). **f**, Low-SNR recording of somatic signals in cortical layer 2/3. **g**, DeepCAD enhanced recording. **h**, Synchronized high-SNR recording (10-fold imaging SNR). Orange arrows point to two neurons. Magnified views of the boxed regions show calcium transients in a ~1.4 s time window. Scale bar 100 μm. **i**, Neurons extracted from the original low-SNR recording (N=150). **j**, Neurons extracted from the DeepCAD enhanced recording (N=229). Manual annotations served as the ground truth. Correctly segmented regions (true positive) are colored green. Missing (false negative) and extra regions (false positive) are colored red and blue, respectively. **k**, Distribution of peak SNRs of extracted calcium traces. CNMF was used for source extraction and peak SNR estimation.

In summary, we demonstrate DeepCAD, a deep self-supervised learning-based method for spatiotemporal enhancement of calcium imaging. Quantitative evaluation on both simulated and experimental data shows that the accuracy of neuron extraction and spike inference can be largely reinforced after denoising. To fully evaluate the capability and reliability of our method, a customized two-photon microscope was built to capture synchronized low-SNR and high-SNR recordings, which indicates that DeepCAD enables a more than tenfold improvement in imaging SNR. To maximize its accessibility, we released an open-source Fiji plugin (Supplementary Fig. 16 and Supplementary Notes 3) and a pre-trained DeepCAD model for two-photon imaging of neuron populations. Our method can be efficiently configured on a common desktop and achieve comparable performance on different imaging systems regardless of objectives and detectors (Supplementary Fig. 17 and Supplementary Video 4). Although DeepCAD is currently investigated only on two-photon microscopy, it can be easily extended to other imaging modalities such as wide-field microscopy and light-sheet microscopy. We anticipate that this method could serve as a general processing step for calcium imaging in photon-limited conditions and promote long-term and high-fidelity recording of neural activities.

## Methods

### Optical setup

A two-photon imaging system was designed to capture strictly synchronized low-SNR and high-SNR calcium recordings for validation of our method. Our system was based on a standard two-photon laser scanning microscope (2PLSM) and the detection path was specially designed to split the fluorescence in a ratio of 1:10. All components of our imaging system are commercially available or easy to fabricate. The schematic of the custom-built two-photon microscope is shown in Supplementary Fig. 12. At the forefront of the optical path, a titanium-sapphire laser system with tunable wavelength (Mai Tai HP, Spectra-Physics) was used as the illumination source to emit the linearly polarized, femtosecond-pulsed Gaussian excitation beam (920 nm central wavelength, pulse width <100 fs, 80 MHz repetition rate). A half-wave plate (AQWP10M-980, Thorlabs) was used to adjust the polarization of the laser beam. Then the laser beam went through an electro-optic modulator (350-80LA-02, Conoptics) to modulate the excitation power and the half-wave plate was rotated to make the electro-optic modulator have maximal extinction ratio. A 4f system composed of two achromatic lenses (AC508-200-B, Thorlabs) with the same focal length was followed to collimate the laser beam. Another 4f system (AC508-100-B and AC508-400-B, Thorlabs) with a fourfold magnification was used to expand the laser beam and guide the beam into a galvo-resonant scanner (8315K/CRS8K, Cambridge Technology) for fast optical scanning. The scanner mount was optimally designed for reliable and distortion-free scanning. Then the beam went through a scan lens (SL50-2P2, Thorlabs) and a tube lens (TTL200MP, Thorlabs) and converged into a tight focus through a high numerical aperture (NA) water-dipping objective (25×/1.05 NA, XLPLN25XWMP2, Olympus). A high-precision piezo actuator (P-725, Physik Instrumente) was additionally used to drive the objective for fast axial scanning. The beam size at the back aperture of the objective was further restricted with an iris set behind the beam expander (L4) to keep the back aperture of the objective underfilled. The effective excitation NA was about 0.5 in our imaging experiments.

For the detection path, fluorescence excited by the Gaussian focus was first collected by the objective. High-NA detection is helpful to detect more fluorescence photons and improve the signal intensity. A long-pass dichroic mirror (DMLP650L, Thorlabs) was used to separate fluorescence by reflecting the fluorescence signals and transmitting the excitation light. A 1:9 (reflectance: Transmission) non-polarizing plate beam splitter (BSN10, Thorlabs) was then placed in the detection path. All fluorescence going through the beam splitter will be split into a 10% component (low-SNR path) and a 90% component (high-SNR path), propagating in two orthogonal directions and detected by two photomultiplier tubes (PMT1001, Thorlabs). A pair of fluorescence filters (MF525-39, Thorlabs; ET510/80M, Chroma) was configured in front of each PMT to fully block wavelengths outside the emission passband of green fluorescent protein (GFP). To improve detection efficiency, we conjugated the back aperture of the objective to the sensor planes of the two PMTs using two 4f systems (TTL200-A and AC254-050-A, Thorlabs). The two detection paths recorded synchronized fluorescence signals but with quite different imaging SNR. Although the high-SNR recording still suffers from noise, it can be used as the reference to identify underlying structures and calcium fluctuations. The field-of-view (FOV) of our two-photon imaging system is about 600 μm and the frame rate is about 30 Hz.

### System calibration

To confirm the fluorescence intensity ratio between the high-SNR detection path and the low-SNR detection path, we imaged 1 μm green-fluorescent beads (G0100, ThermoFisher) for system calibration. The beads suspension was first diluted and embedded in 1.0% agarose and then mounted on a microscope slide to form a single beads layer composed of sparse beads. A specified region was continuously scanned to acquire 500 consecutive frames. These frames can be regarded as independent samplings of the same underlying scene. To reduce the impact of detection noise, we averaged these frames to obtain the noise-free image of each path (Supplementary Fig. 13). All beads in the FOV were manually segmented and the intensity of each bead was calculated by averaging all pixels inside its segmentation mask. According to our statistical analysis, the fluorescence intensity of the high-SNR detection path was approximately tenfold higher than that of the low-SNR detection path.

### Mouse preparation and calcium imaging

All experiments involving mice were performed in accordance with institutional guidelines for animal welfare and have been approved by the Institutional Animal Care and Use Committee (IACUC) of Tsinghua University.

Adult transgenic mice (Ai148D/Rasgrf2-dCre) at 8-12 postnatal weeks were anesthetized with 1.5% isoflurane and craniotomy surgeries were conducted using a stereotaxic instrument (68018, RWD Life Science) under a bright-field binocular microscope (77001S, RWD Life Science). A custom-made coverslip fitting the shape of the cranial window (~6 mm in diameter) was embedded and cemented to the skull. A biocompatible titanium headpost was then cemented to the skull for stable head fixation. The edge of the cranial window was enclosed with dental cement to hold the immersion water of the objective. After the surgery, 0.25mg/g body weight of Trimethoprim (TMP) was intraperitoneally injected to induce the expression of GCaMP6f genetically encoded calcium indicator (GECI) in layer 2/3 neurons across the whole brain. To reduce potential inflammation, 5 mg/kg body weight of Ketoprofen was injected subcutaneously. Each mouse was housed in a separate cage for 1-2 weeks of postoperative recovery.

Imaging experiments were carried out when the cranial window became clear and no inflammation occurred. Mice were first rapidly anesthetized with 3.0% isoflurane and then fixed onto a custom-made holder with the headpost. The mouse holder was mounted on a precision translation stage with three motorized axes (M-VP-25XA-XYZL, Newport) to find the region of interest (ROI) for imaging. The correction ring of the objective was adjusted to compensate for the coverslip thickness and eliminate spherical aberrations. The excitation power after the objective was kept below 140 mW in all experiments to avoid potential laser-induced tissue damage. Gaseous anesthesia was switch off and the mice kept awake during the whole imaging process.

### Network architecture and training details

The network architecture of DeepCAD employs 3D U-Net, which is reported to have superior performance on the segmentation of volumetric data^20^. In general, the network is composed of a 3D encoder module (the contracting path), a 3D decoder module (the expanding path), and three skip connections from the encoder module to the decoder module (Supplementary Fig. 1). In the 3D encoder module, there are three encoder blocks. Each block consists of two 3×3×3 convolutional layers followed by a leaky rectified linear unit (LeakyReLU) and a 2×2×2 max pooling with strides of two in three dimensions. In the decoder module, there are three decoder blocks, each of which contains two 3×3×3 convolutional layers followed by a LeakyReLU and a 3D nearest interpolation. A group normalization^22^ layer is configured after each convolutional layer. The skip connections link low-level features and high-level features by concatenating their feature maps. All operations (convolutions, max poolings, and interpolations) in the network are in 3D to aggregate spatial information and temporal information. For the loss function, we used the arithmetic average of a L1-norm loss term and a L2-norm loss term. The model was trained on 3D tiles with a spatial size of 64×64 pixels and a temporal size of 300 frames. Small spatial size can lower memory requirements and reduce the training time, and large temporal size is helpful to make full use of temporal information.

Adam optimizer^23^ was used for network training with a learning rate of 0.00005 and exponential decay rates of 0.5 for the first moment and 0.9 for the second moment. We used Graphics Processing Units (GPU) to accelerate the training and test process. It took about 12 hours to train our model for 20 epochs on a typical training set (about 1200 3D tiles) with a single GPU (Nvidia TITAN RTX, 24 GB memory). Training time can be further shortened by using a more powerful GPU or parallelizing the training process on multiple GPUs.

The full 3D architecture of DeepCAD makes it easy to overfit because 3D convolutions usually involve more parameters than the 2D counterpart. The best denoising performance is only achieved at the point where there is neither underfitting nor overfitting. To screen out the model with the best generalization ability, we saved the network snapshot after each training epoch and evaluated its performance on a holdout validation set. We fed the validation data into each model and calculated the standard deviation projection of the output stack of each model. Then, the average pixel intensity was calculated on a small dark region (*e.g*. blood vessels or a small region without neural activity during the recording) of all standard deviation projections. The best model was selected to be the one with the smallest dark standard deviation.

### Data simulation

Our simulation program includes a step for synthesizing the noise-free video (ground truth) and a step for adding the Mixed Poisson-Gaussian (MPG) noise (Supplementary Notes 1-2). Firstly, to generate realistic simulated calcium imaging data, we constructed a neuron library containing the spatial profiles of 517 neurons. These neurons were extracted using the constrained nonnegative matrix factorization algorithm^21^ (CNMF) from an experimentally obtained two-photon calcium imaging data of a virus-transfected wild-type mouse expressing GCaMP6f (layer 2/3 at the primary somatosensory cortex). For the spatial component that defines the location of each neuron, 120 neurons were randomly selected from the library to keep the sparsity of neurons. For the temporal component that defines the fluorescence fluctuations of each neuron, MLspike^24^ was employed to generate calcium traces with GCaMP6f kinetics. Then, these two components were reshaped into 2D matrices and the simulated noise-free data (1 μm/pixel spatial sampling rate, 30 Hz frame rate) was synthesized as the product of the spatial matrix and the temporal matrix. The noise-contaminated counterpart was ultimately generated by adding the content-related MPG noise. Data with different imaging SNRs were simulated with different relative photon numbers. Their relationship was investigated in Supplementary Fig. 7. All images were saved as uncompressed tif files with the format of unsigned 16-bit integer (uint16). More details of data simulation and related mathematical models are described in Supplementary Notes 1-2.

### Single-neuron recordings

The data of simultaneous two-photon imaging and electrophysiological recordings of single-neuron activities were released by the Svoboda lab^25^ and were downloaded from the Collaborative Research in Computational Neuroscience (CRCNS) platform. Only recordings of GCaMP6f neurons were used in this study. The image stacks were fourfold downsampled to reduce the sampling rate and some outlier recordings with very sparse spikes and low electrophysiological SNR were excluded. Fluorescence traces were extracted from temporal stacks using manually annotated cytoplasmic masks. For spike inference, we used the MLspike algorithm^24^, which was reported to rank first in the *Spikefinder* challenge^26^. All traces were divided by their mean values for normalization before fed into the spike inference pipeline. Recommended model parameters for GCaMP6f indicator were used to ensure optimal performance of spike inference.

### Data analysis of neuronal populations

Calcium imaging data of large neuronal populations were first registered with a non-rigid motion correction method^27^ and the black edges of registered images were clipped. Then, CNMF^21^ was employed as the source extraction method for neuron segmentation and trace extraction. A spatial matrix and a temporal matrix can be obtained from each video, storing the spatial footprints and corresponding calcium traces of all active neurons, respectively. The same set of parameters was used for the original low-SNR recording and corresponding DeepCAD enhanced counterpart, as well as the high-SNR recording. Simulated data were analyzed following the same pipeline except motion correction. Along with automatic neuron extraction, we also performed manual annotations to inspect our results. High-SNR recordings were tenfold downsampled along the time axis by averaging each consecutive ten frames, which reduced the disturbance of detection noise and was helpful to improve annotation accuracy. Boundaries of all active components were annotated using the ROI Manager toolbox of Fiji. The final segmentation masks were generated through subsequent morphological operations of images and connected domain extraction implemented with customized MATLAB scripts.

### Performance metrics

Two types of metrics were used for quantitative evaluation of the spatial and temporal performance of DeepCAD. For synthetic calcium imaging data, corresponding clean images and ground-truth calcium traces were available. SNR and PSNR were used as the spatial metric to evaluate pixel-level similarity between DeepCAD enhanced images and ground-truth images. Pearson correlation coefficient (R) was used as the temporal metric to reflect the similarity between enhanced traces and ground-truth traces. The Pearson correlation between signal *x* and the reference signal *y* is defined as

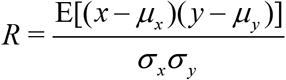

where *μ_x_* and *μ_y_* are the mean values of signal *x* and *y*, respectively; *σ_x_* and *σ_y_* are the standard deviations of signal *x* and *y*, respectively; E represents arithmetic mean.

Furthermore, we also evaluated the performance of DeepCAD based on more complex downstream tasks such as neuron extraction and spike inference, which are the most crucial prerequisites in functional analysis of neural circuits from calcium imaging data. We considered neuron extraction as an instance segmentation problem and adopted an object-level metric to evaluate segmentation performance^28^. Different intersection-over-union (IoU, defined as the intersection area divided by the union area of two objects) thresholds were selected to determine correctly segmented objects. For a specified IoU threshold, the segmentation accuracy (F1 score) was defined as the harmonic mean of sensitivity and precision, which can be formulated as

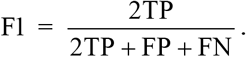

Here, TP, FP, and FN are the number of true positives, false positives, and false negatives, respectively. When applied CNMF as the source extraction method, the SNR of calcium traces was quantified with the peak SNR automatically calculated by the CaImAn toolbox^29^ with infinite outliers eliminated. For spike inference, we used the error rate (ER) to quantify the performance of spike inference, which is defined as ER = 1-F1. Spikes detected from simultaneous electrophysiological recordings were used as the ground truth for ER calculation. The evaluation process was implemented with customized MATLAB scripts. SNR, PSNR, Pearson correlation coefficient, and IoU were computed using built-in functions.

## Data availability

Our data will be made publicly available post peer-review.

## Code availability

Our python code and Fiji plugin will be made publicly available post peer-review.

## Acknowledgements

We would like to acknowledge Y. Tang and Y. Yang at the School of Medicine of Tsinghua University for providing transgenic mice for imaging and the mesoscope imaging data for cross-system validation. We thank the Svoboda lab at Janelia Research Campus for releasing their data of simultaneous electrophysiology and two-photon recording. This work was supported by the National Natural Science Foundation of China (62088102, 61831014, 61531014 and 6181001011) and the Shenzhen Science and Technology Project under Grant (ZDYBH201900000002 and JCYJ20180508152042002).

## Author Contributions

Q. D., H. W., L. F. and XY. L. conceived this project. Q. D., H. W. and L. F. supervised this research. XY. L. and G. Z. designed detailed implementations and processed the data. XY. L designed and set up the imaging system. XY. L and G. Z. conducted the experiments. G. Z. developed the python code and the Fiji plugin. J. W., Y. Z., and X. L. directed the experiments and data analysis. L. F., Y. Z., Z. Z, H. Q. and H. X. gave critical support on system setup and imaging procedure. J. W., L. F., Y. Z., X. L., H. Q., H. X., H. W. and Q. D. gave critical discussions on the results. All authors participated in the writing of the paper.

