## Supplementary Information for "Reinforcing neuron extraction and spike inference in calcium imaging using deep self-supervised learning"

**Supplementary Information Table**

| | |
|---|---|
| **Supplementary Figure 1** | Network architecture. |
| **Supplementary Figure 2** | Comparison of denoising methods. |
| **Supplementary Figure 3** | Data processing pipeline. |
| **Supplementary Figure 4** | Interpretability of DeepCAD model. |
| **Supplementary Figure 5** | Temporal scalability of DeepCAD. |
| **Supplementary Figure 6** | Spatial scalability of DeepCAD. |
| **Supplementary Figure 7** | Properties of simulated calcium imaging data. |
| **Supplementary Figure 8** | Denoising performance on different noise levels. |
| **Supplementary Figure 9** | Calcium traces before and after denoising. |
| **Supplementary Figure 10** | Single-pixel fluorescence. |
| **Supplementary Figure 11** | DeepCAD reduces the error rate of spike inference under different input SNRs. |
| **Supplementary Figure 12** | Simultaneous low-SNR and high-SNR two-photon imaging system. |
| **Supplementary Figure 13** | System calibration. |
| **Supplementary Figure 14** | Pixels for extracting single-pixel fluorescence. |
| **Supplementary Figure 15** | Human inspection of segmentation results. |
| **Supplementary Figure 16** | Fiji-based DeepCAD plugin. |
| **Supplementary Figure 17** | Cross-system validation. |
| **Supplementary Note 1** | Mixed Poisson-Gaussian Noise. |
| **Supplementary Note 2** | Generation of simulated calcium imaging data |
| **Supplementary Note 3** | Tutorial of DeepCAD plugin |

#### Supplementary Videos

| | |
|---|---|
| **Supplementary Video 1** | Denoising performance of DeepCAD on single neuron recordings. The top panel shows simultaneous electrophysiological recording of the neuron, which reflects the ground-truth neural activity. Detected spikes are marked with black dots. The original noisy data and DeepCAD enhanced data are shown in the middle panel and the bottom panel, respectively. |
| **Supplementary Video 2** | DeepCAD massively enhanced the recording of neuropil activities in cortical layer 1 of a mouse expressing GCaMP6f. The low-SNR recording, DeepCAD enhanced recording, and the high-SNR recording are played synchronously. Below are magnified views of the boxed regions. The bottom panel shows fluorescence traces extracted from 40 dendritic pixels. |
| **Supplementary Video 3** | From left to right are the low-SNR recording of spontaneous calcium transients of a large neuronal population (layer 2/3, GCaMP6f), the DeepCAD enhanced counterpart, and corresponding high-SNR recording, respectively. The bottom panel shows magnified views of boxed regions. |
| **Supplementary Video 4** | Denoising performance of DeepCAD on three two-photon laser-scanning microscopes (2PLSMs) with different system setups. Our system was equipped with alkali PMTs (PMT1001, Thorlabs) and a 25×/1.05 NA commercial objective (XLPLN25XWMP2, Olympus). The standard 2PLSM was equipped with a GaAsP PMT (H10770PA-40, Hamamatsu) and a 25×/1.05 NA commercial objective (XLPLN25XWMP2, Olympus). The two-photon mesoscope was equipped with a GaAsP PMT (H11706-40, Hamamatsu) and a 2.3×/0.6 NA custom objective. The same pre-trained model was used for denoising these data. |

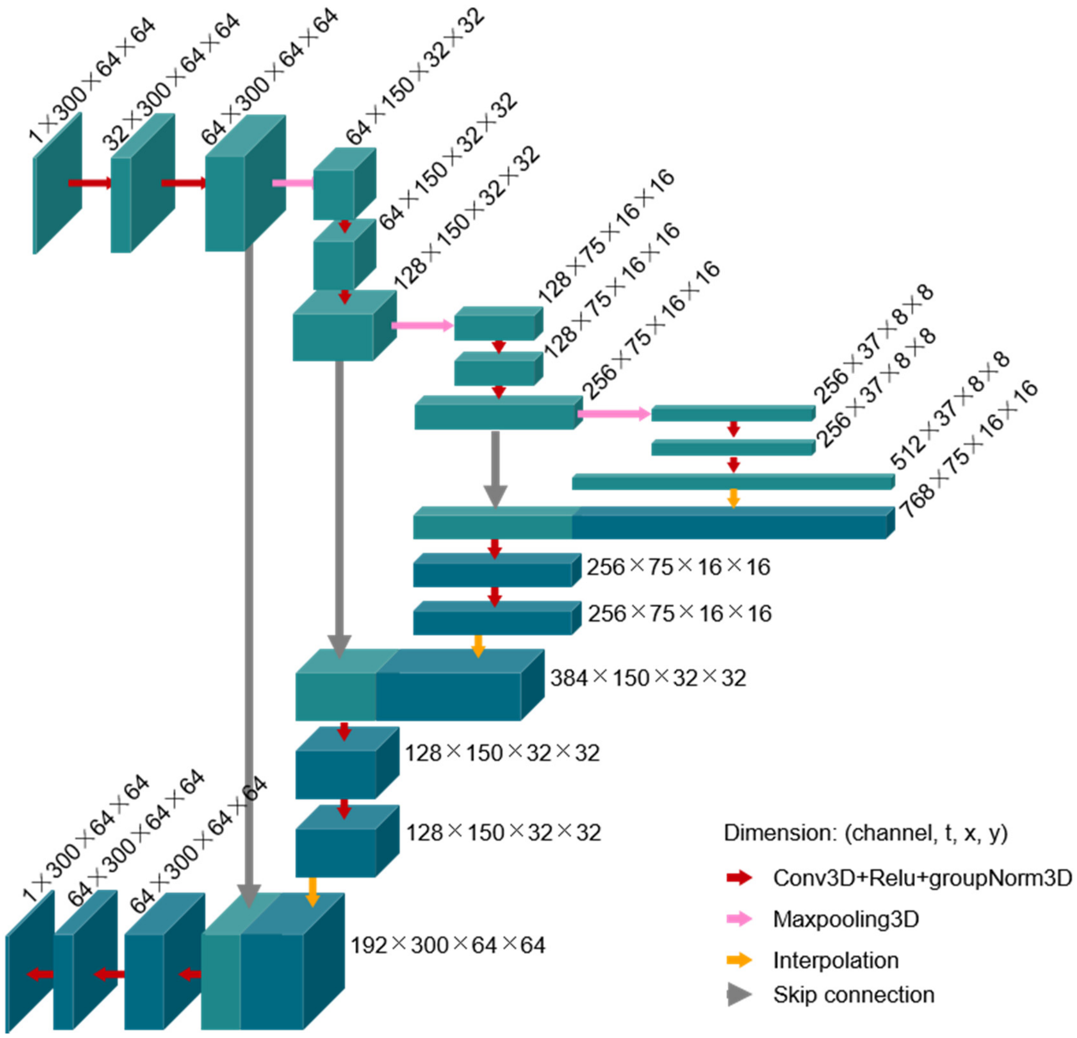


**Supplementary Figure 1**

**Network architecture.**

Our model adopted 3D U-net[1], which is composed of a 3D encoder module, a 3D decoder module, and three skip connections from the encoder module to the decoder module. In the encoder module, there are three encoder blocks. Each block consists of two 3×3×3 convolutional layers followed by a leaky rectified linear unit (LeakyReLU), a group normalization layer, a 2×2×2 max pooling with strides of 2 in three dimensions. In the decoder module, there are three decoder blocks, each of which contains two 3×3×3 convolutional layers followed by a LeakyReLU, a group normalization layer, and a 3D nearest interpolation. The skip connections pass feature maps from the encoder module to the decoder module to integrate low-level features and high-level features. Feature maps of the encoder module and the decoder module are represented in different colors. All operations are in 3D and feature maps are all 4D tensors. 3D feature maps were used here to simplify representation.

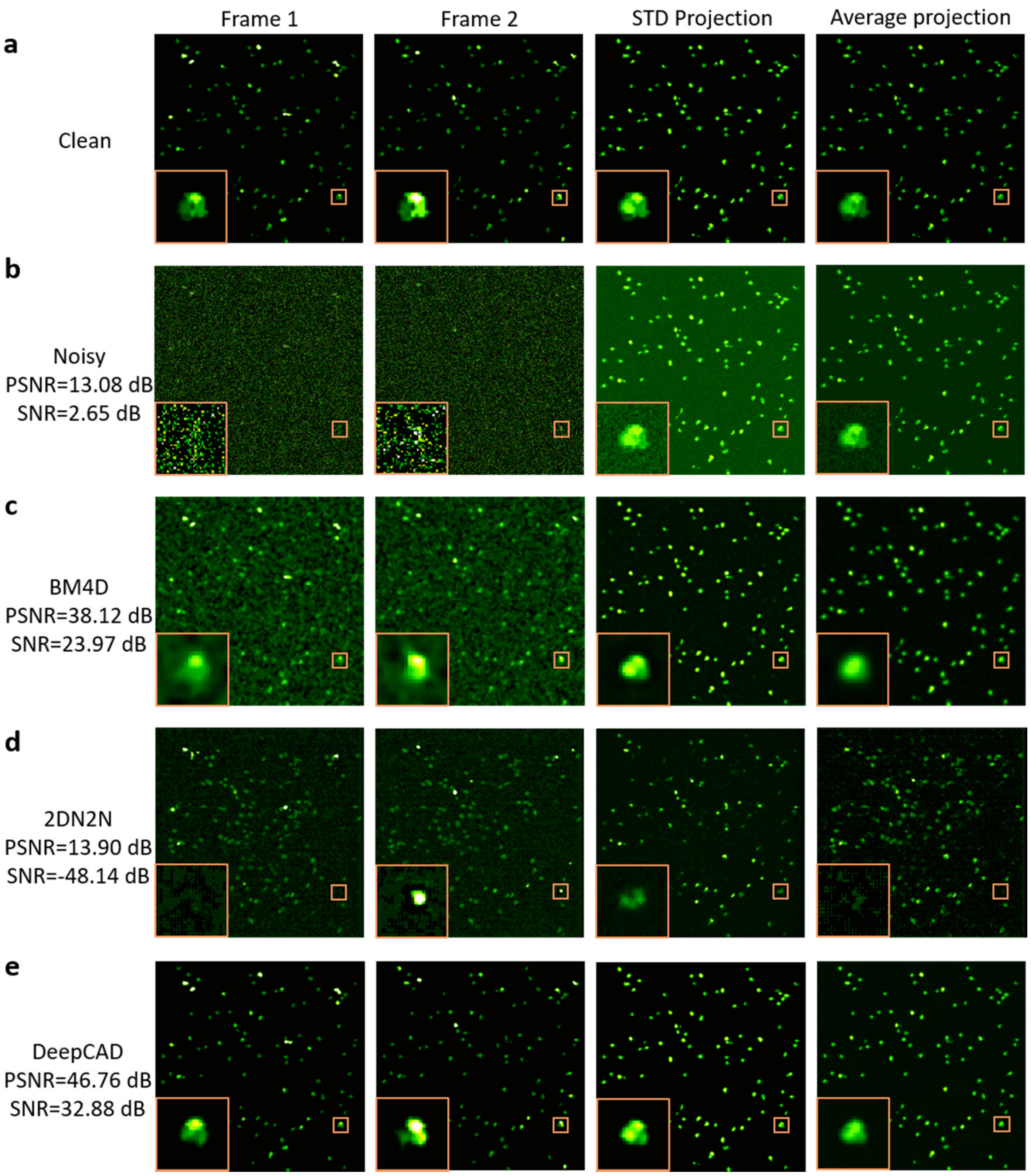

Frame 1
Frame 2
STD Projection
Average projection
a
Clean
b
Noisy
PSNR=13.08 dB
SNR=2.65 dB
c
BM4D
PSNR=38.12 dB
SNR=23.97 dB
d
2DN2N
PSNR=13.90 dB
SNR=-48.14 dB
e
DeepCAD
PSNR=46.76 dB
SNR=32.88 dB

**Supplementary Figure 2**

**Comparison of denoising methods.**

Three denoising methods are demonstrated here, including BM4D[2], 2D noise2noise[3] (2DN2N), and DeepCAD. 2DN2N was based on 2D U-net[4] and trained with image pairs composed of two consecutive frames (one as the input and the other as the target). BM4D is the extension of BM3D[5] for denoising of volumetric data. This experiment was carried out on synthetic calcium imaging data (Supplementary Note 1-2) because corresponding ground-truth images are accessible for quantitative evaluation. From the leftmost column to the rightmost column are two frames at two moments, standard deviation (STD) projections, and average projections. Magnified views of three overlapping neurons are shown at the left bottom of each image. **a**, Clean data without detection noise. **b**, Raw data before denoising. **c**, Results of BM4D. Although PSNR has been improved a lot, noise cannot be effectively removed and there appeared random speckles and images are also blurred. **d**, Results of 2DN2N. Caused by the absence of temporal information in its 2D architecture, 2DN2N cannot determine whether there's a neuron or not and would produce artifacts. **e.** Results of DeepCAD. Accurate neuron location and fluorescence restoration can be achieved because of the 3D architecture of DeepCAD. Signal-to-noise ratios (SNRs) and Peak signal-to-noise ratios (PSNRs) were averaged on all 3600 frames of the stack.

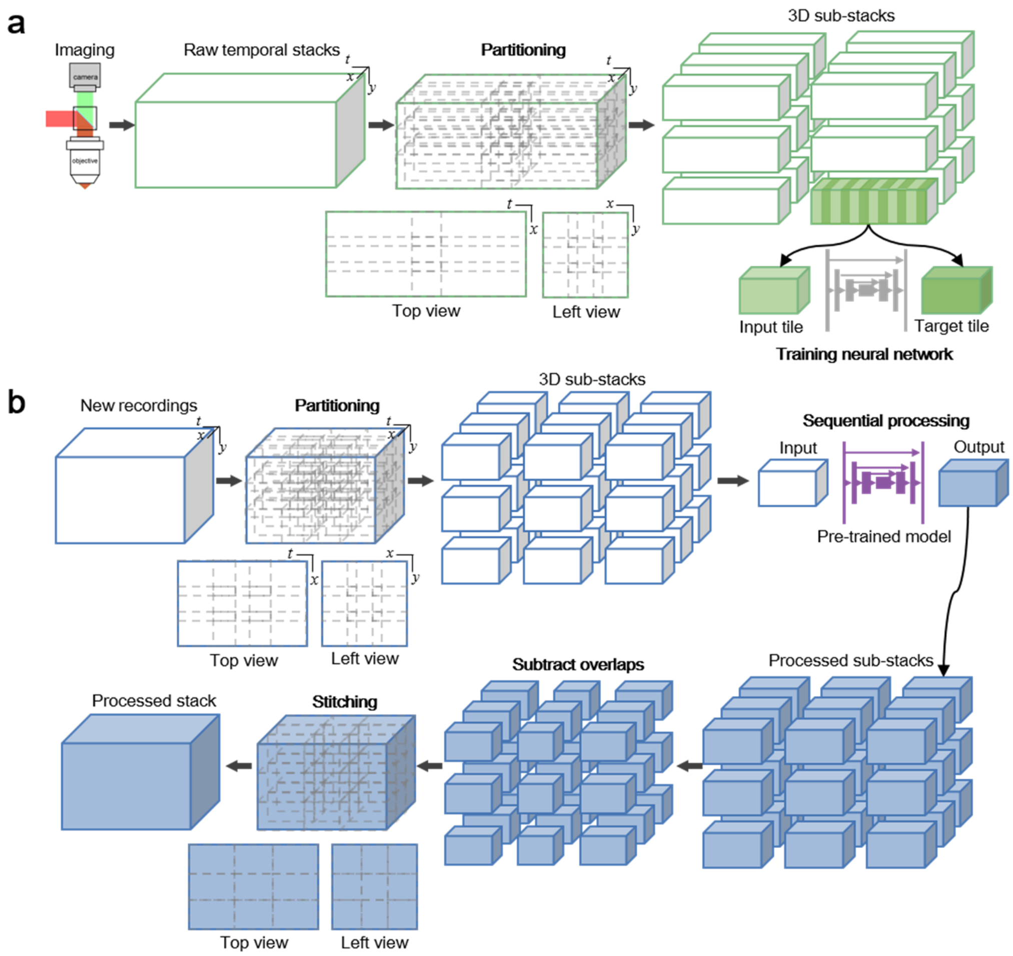


**Supplementary Figure 3**

**Data processing pipeline.**

**a**, The training process. Raw data captured by the imaging system is organized in a 3D form ($x$, $y$, $t$) and saved as a temporal stack. The original stack was partitioned into thousands of 3D sub-stacks (64×64×600 pixels) with about 25% overlap in each dimension. For temporal stacks with a small lateral size or short recording period, sub-stacks can be randomly cropped from the original stack to augment the training set. Then, interlaced frames of each sub-stack are extracted to form two 3D tiles (64×64×300 pixels). One of them serves as the input and the other serves as the target for network training. **b**, Application of the pre-trained model. New recordings obtained with the imaging system are partitioned into 3D sub-stacks (64×64×300 pixels) with 25% overlap in each dimension. Then, pre-trained models saved during the training process are loaded into memory and the sub-stacks are directly fed into the model. Enhanced sub-stacks are sequentially output from the network and overlapping regions (both the lateral and temporal overlaps) are subtracted from the output sub-stacks. The final enhanced stack can be obtained by stitching all 3D sub-stacks.

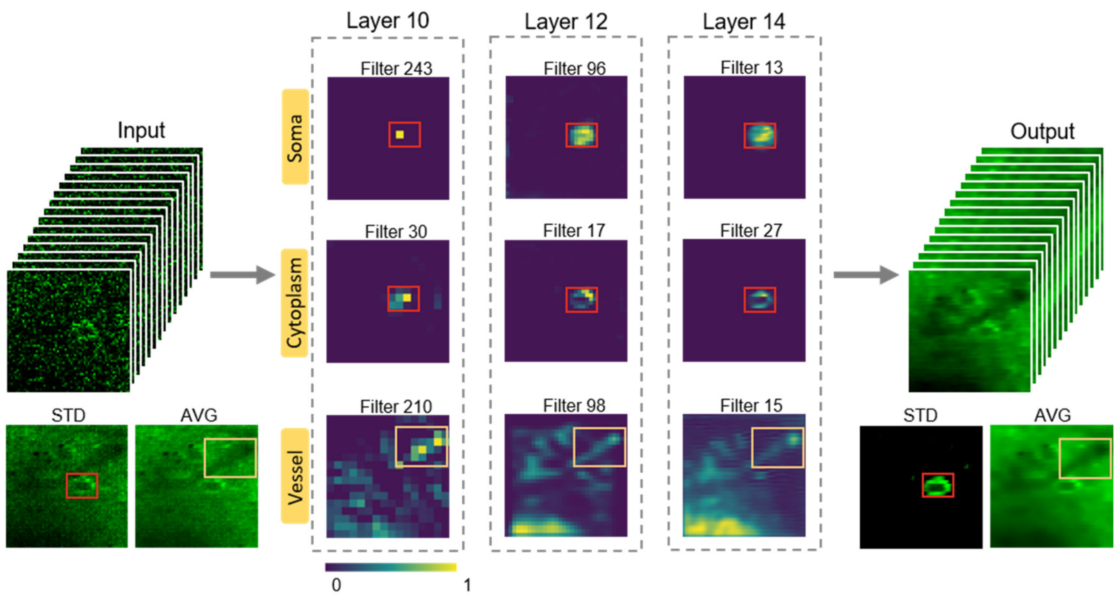


**Supplementary Figure 4**

**Interpretability of DeepCAD model.**

To demonstrate the interpretability and reliability of our pre-trained DeepCAD model, A small 3D patch (64×64×300 pixels) was input into the model and feature maps of convolutional layers were visualized[6]. Typical feature maps of three intermediate convolutional layers in the decoder module (Layer 10, Layer 12, and Layer 14) are shown here, displayed as the average intensity projection (AVG) of original 3D feature maps. The feature representations learned by DeepCAD have significant semantic meaning, such as soma-like structures, cytoplasm-like structures, and vessel-like structures (or shadows). These interpretable semantic representations would contribute to locating neurons, restoring cytoplasmic fluorescence, and avoiding unwanted intensity fluctuations in shadow areas.

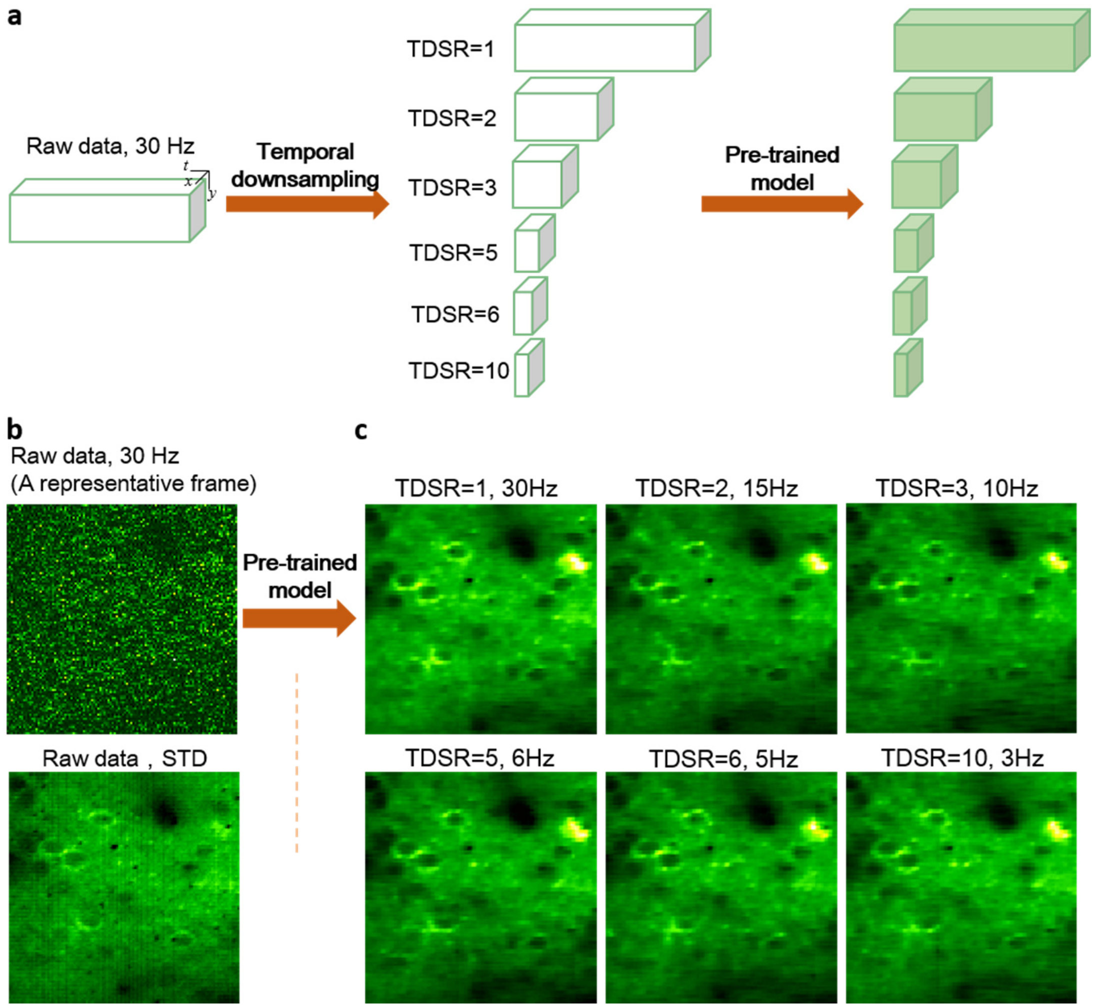


**Supplementary Figure 5**

**Temporal scalability of DeepCAD.**

**a**, Raw data (3900 frames, 30 Hz frame rate) obtained with a two-photon microscope at low excitation dosage was used to train a DeepCAD model for noise removal. Stacks imaged at various frame rates were obtained through downsampling the original stack at different temporal downsampled rates (TDSR=1, 2, 3, 5, 6, 10). Then the downsampled stacks with lower imaging speed were fed into the pre-trained model and corresponding enhanced stacks were output from the model. **b**, Original stacks before the enhancement of DeepCAD were severely corrupted in noise. A representative frame is shown here to indicate the noise level. The standard deviation projection (STD) of the original stack is also shown to indicate locations of neurons and vessels. **c,** One common frame of all downsampled stacks after enhancement of the pre-trained model. Although imaged at different frame rates, DeepCAD has non-inferior performance on a wide range of temporal resolution.

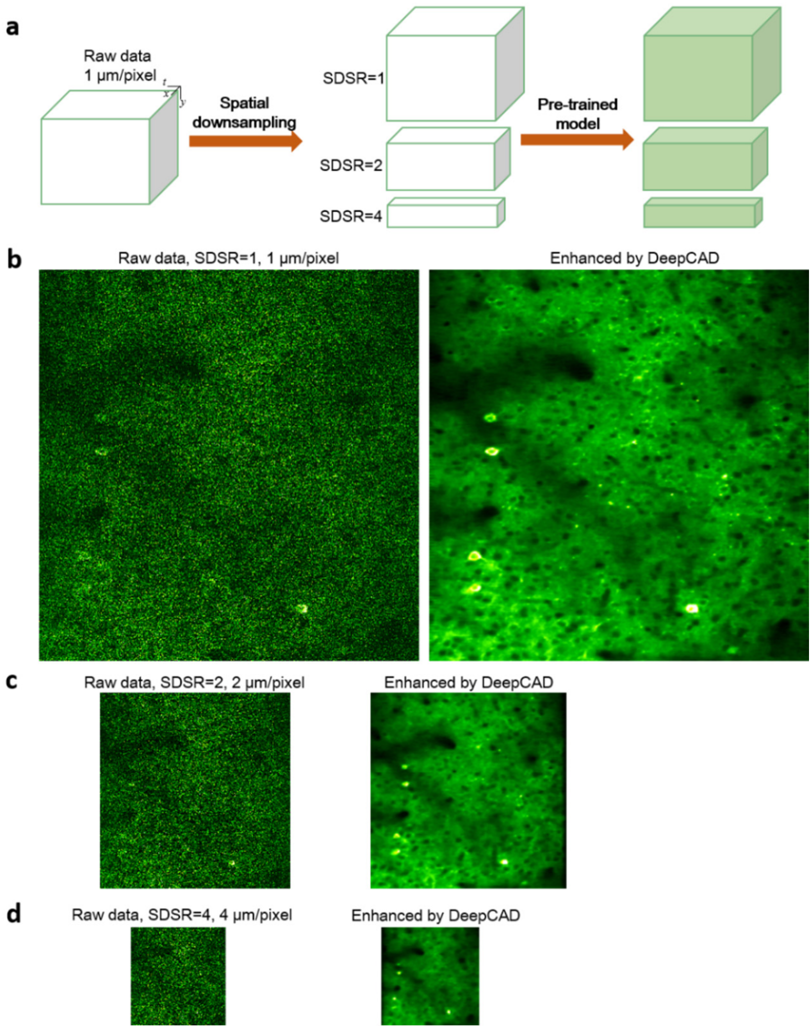


**Supplementary Figure 6**

**Spatial scalability of DeepCAD.**

Raw data (3900 frames, 1 μm/pixel) obtained with a two-photon microscope at low excitation dosage was used to train a DeepCAD model for noise removal. Stacks imaged at various spatial sampling rate were obtained through downsampling (without interpolation) the original stack at different spatial downsampled rates (SDSR=1, 2, 4). Then the downsampled stacks with lower spatial resolution were fed into the pre-trained model and corresponding enhanced stacks were output from the model. **b**, The original stack (SDSR=1, 1 μm/pixel) before and after the enhancement of the pre-trained model. **c**, The twofold spatial downsampled stack (SDSR=2, 2 μm/pixel) before and after the enhancement of the pre-trained model. **d**, The fourfold spatial downsampled stack (SDSR=4, 4 μm/pixel) before and after the enhancement of the pre-trained model. Only one representative frame is shown here. Similar performance was achieved on all other frames (~3900). Although imaged at different spatial sampling rates, DeepCAD has non-inferior performance on a wide range of spatial resolution.

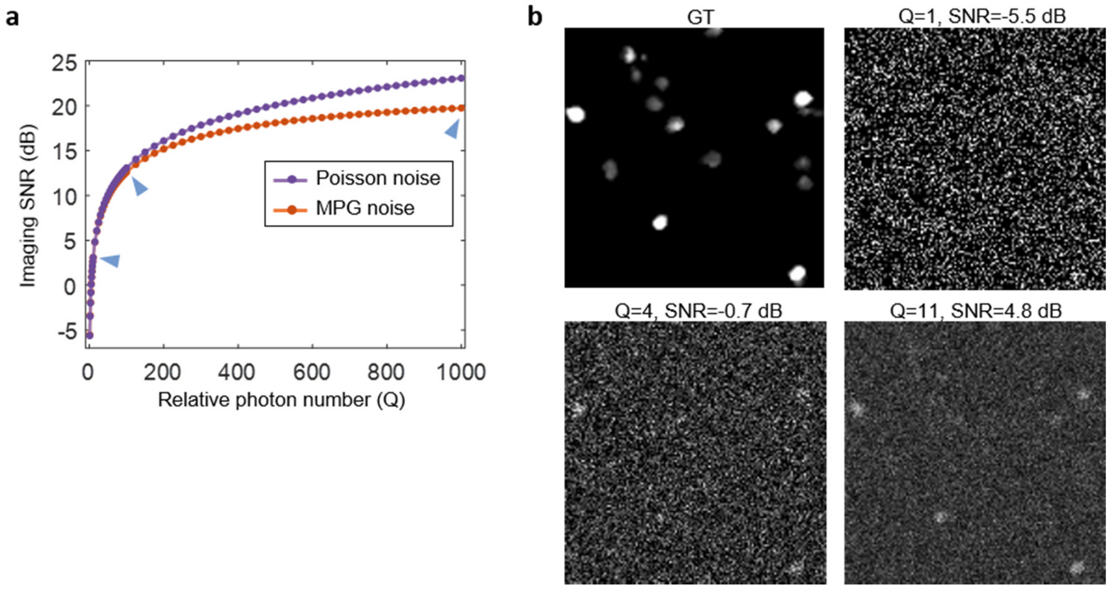


**Supplementary Figure 7**

**Properties of simulated calcium imaging data.**

**a,** The relation between imaging SNR and relative photon number. Both Poisson noise and Mixed Poisson-Gaussian (MPG) noise are shown. **b**, Example images with MPG noise of different imaging SNR.

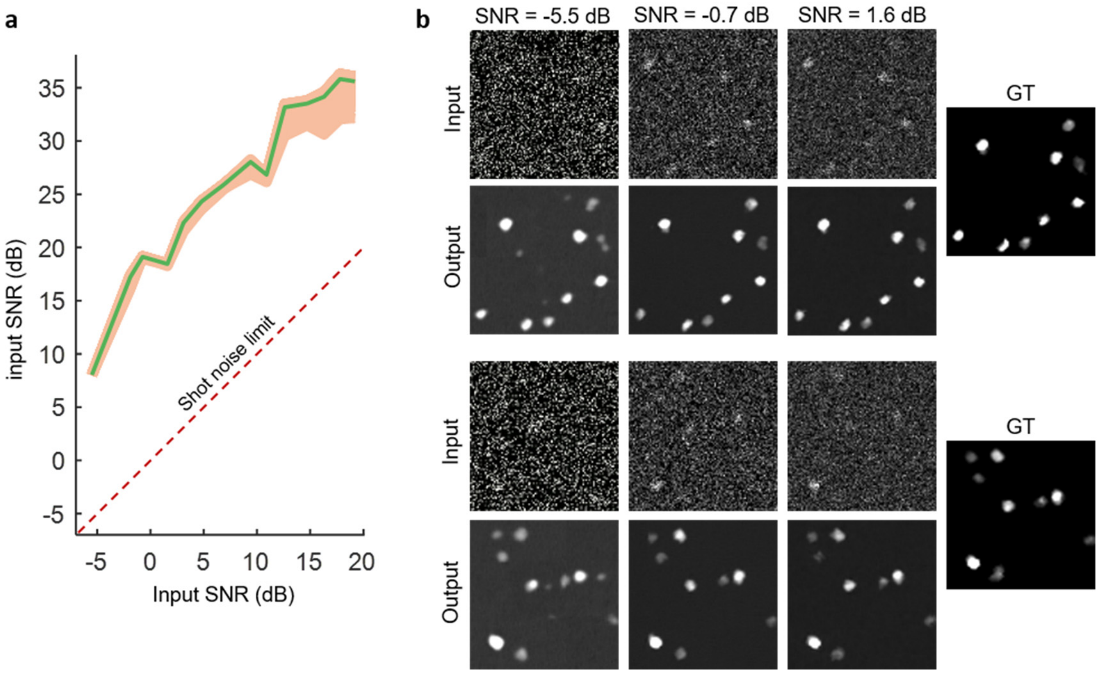


**Supplementary Figure 8**

**Denoising performance on different noise levels.**

Simulated calcium imaging stacks (3600 frames, 30 Hz frame rate, containing 120 neurons) with different noise levels (quantified with peak signal-to-noise ratios (PSNR)) were generated using our synthetic model (Supplementary Note 1-2). Then, DeepCAD models were trained for denoising of these stacks. **a**, The relationship between SNR of input images and that of output images. Green line represents average SNR of all images and the yellow region is the range of all values. Significant improvements were observed across different input SNRs, about 18 dB on average. The red dashed line represents the shot noise limit, *i.e.*, the input SNR is equal to the output SNR. **b**, Examples of denoising performance on different input SNRs. Slight artifacts would appear if the input SNR is very low (≤-5.5 dB).

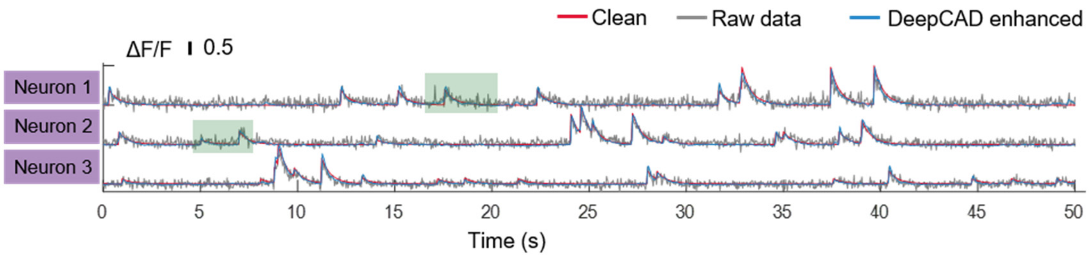


**Supplementary Figure 9**

**Calcium traces before and after denoising.**

Calcium traces of three neurons were extracted from the clean data (red), the raw noisy data (gray), and the DeepCAD enhanced counterpart (blue). Neuron profiles for data synthesis were used as the segmentation masks. Magnified views of green shaded regions were shown in Fig. 1g.

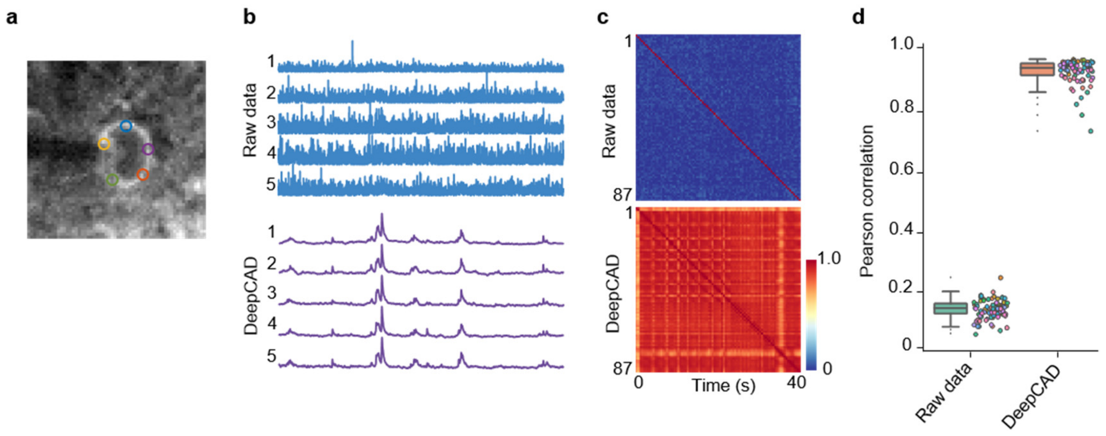


**Supplementary Figure 10**

**Single-pixel fluorescence.**

**a**, Five pixels for extracting sigle-pixel fluorescence. **b**, Fluorescence traces of five cytoplasmic pixels before (top) and after (bottom) denoising. **c**, Cross correlation matrices of all 87 cytoplasmic pixels before (top) and after (bottom) denoising. The denoised traces had strong correlation because the random detection noise was effectively suppressed. **d**, Pearson correlation coefficients between single-pixel fluorescence (N=87) and whole-cytoplasm fluorescence (the average of all pixels in the cytoplasm).

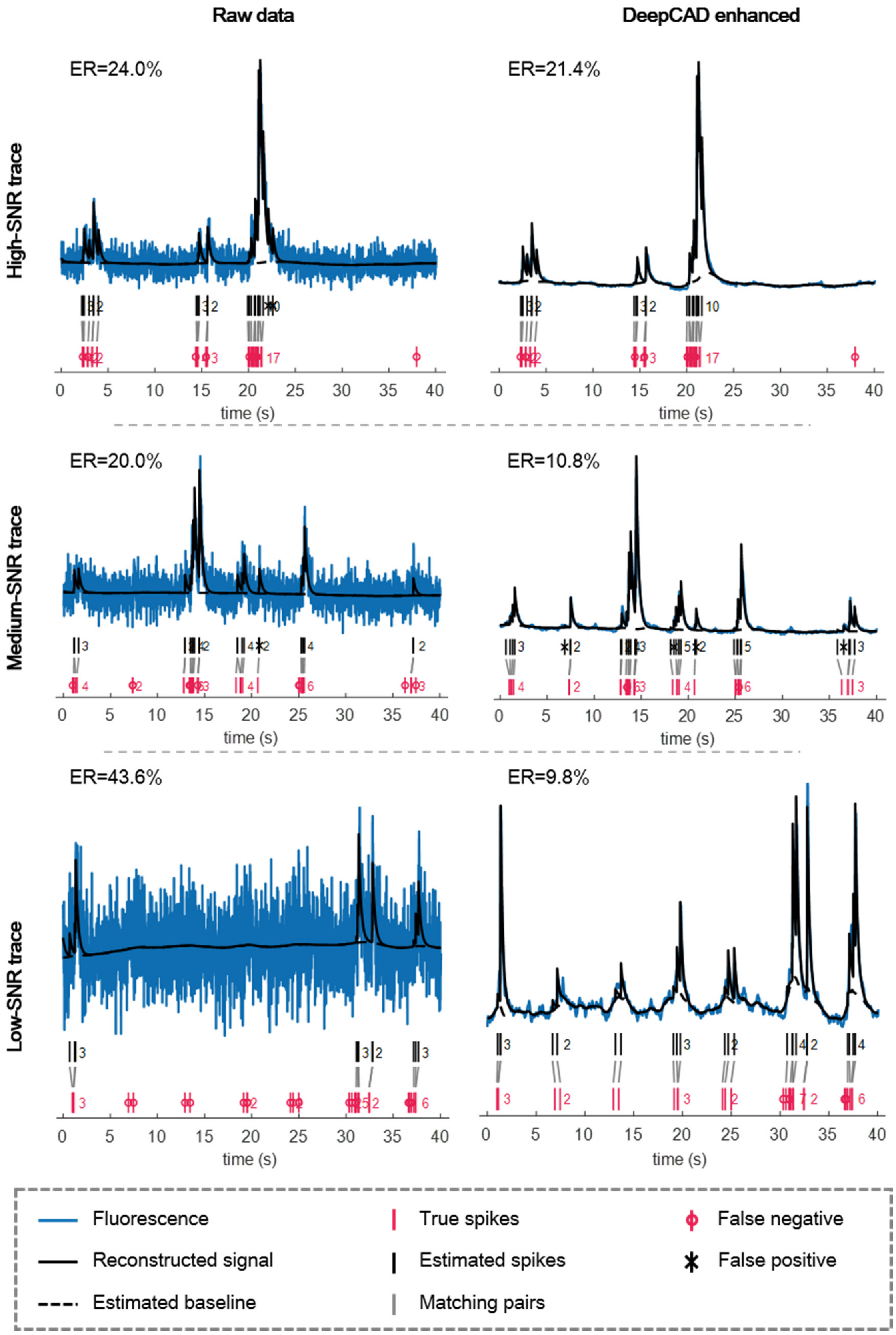

Raw data
DeepCAD enhanced
High-SNR trace
Medium-SNR trace
Low-SNR trace
ER=24.0%
ER=21.4%
ER=20.0%
ER=10.8%
ER=43.6%
ER=9.8%
time (s)
Fluorescence
Reconstructed signal
Estimated baseline
True spikes
Estimated spikes
Matching pairs
False negative
False positive

**Supplementary Figure 11**

**DeepCAD reduces the error rate of spike inference under different input SNRs.**

Fluorescence traces and true spike timings were obtained from simultaneous electrophysiology and two-photon imaging of neurons expressing GCaMP6f[7,8]. The state-of-the-art spike inference algorithm, MLspike[9], which was ranked first in the *Spikefinder* challenge[10], was used to infer spikes from calcium traces. Left column: spike inference of calcium traces extracted from original noisy data. Right column: spike inference of fluorescence traces extracted from DeepCAD enhanced data. Three cases of different SNRs are shown here. From top to bottom are the high-SNR example, the medium-SNR example, and the low-SNR example. For high-SNR recordings, DeepCAD had slight improvement on spike inference. But for low-SNR recordings, significant improvement was observed because of the enhancement of DeepCAD.

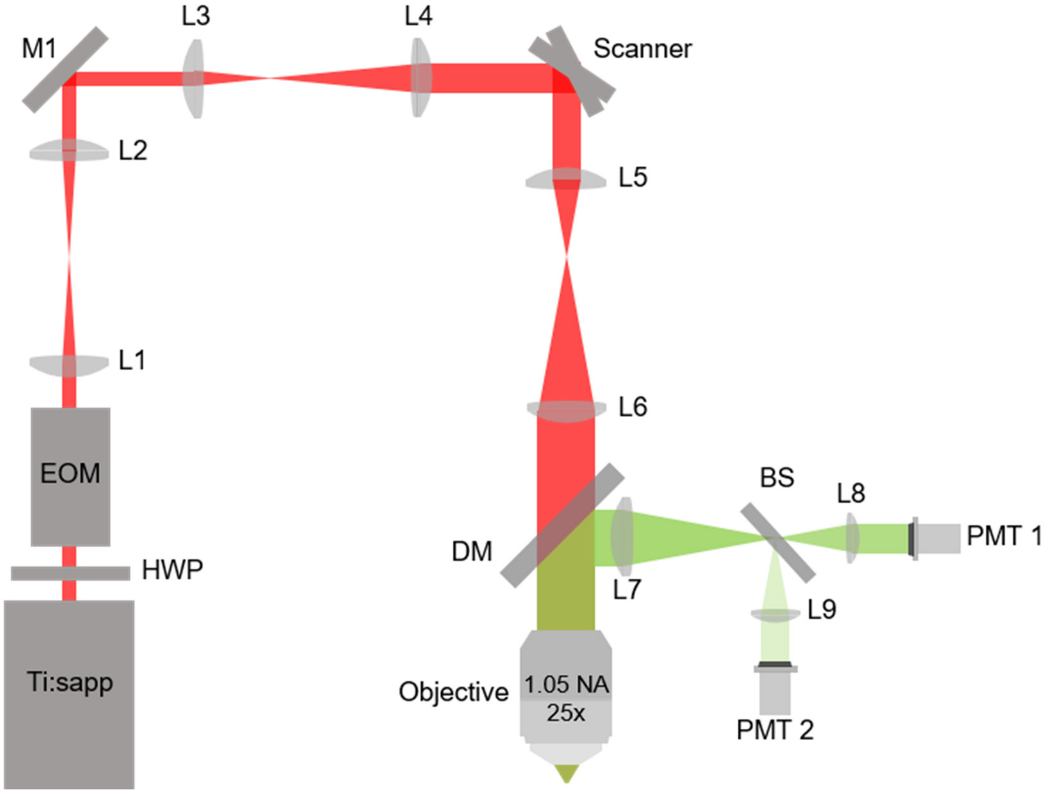


**Supplementary Figure 12**

**Simultaneous low-SNR and high-SNR two-photon imaging system.**

Ti:sapp: titanium-sapphire laser with tunable wavelength; HWP: half-wave plate; EOM: Electro-Optic Modulator; M1: mirror; L1, L2, L3, L4, L5, L6, L7, L8, L9: lens; Scanner: galvo-resonant scanners; DM: long-pass dichroic mirror to separate fluorescence signals (green path) from excitation light (red path); BS: 1:9 (reflectance: transmission) non-polarizing plate beam splitter; PMT1, PMT2: photomultiplier tubes. Fluorescence signals were split into a low-SNR (~10%) component and a high-SNR (~90%) component and were synchronously detected by two PMTs.

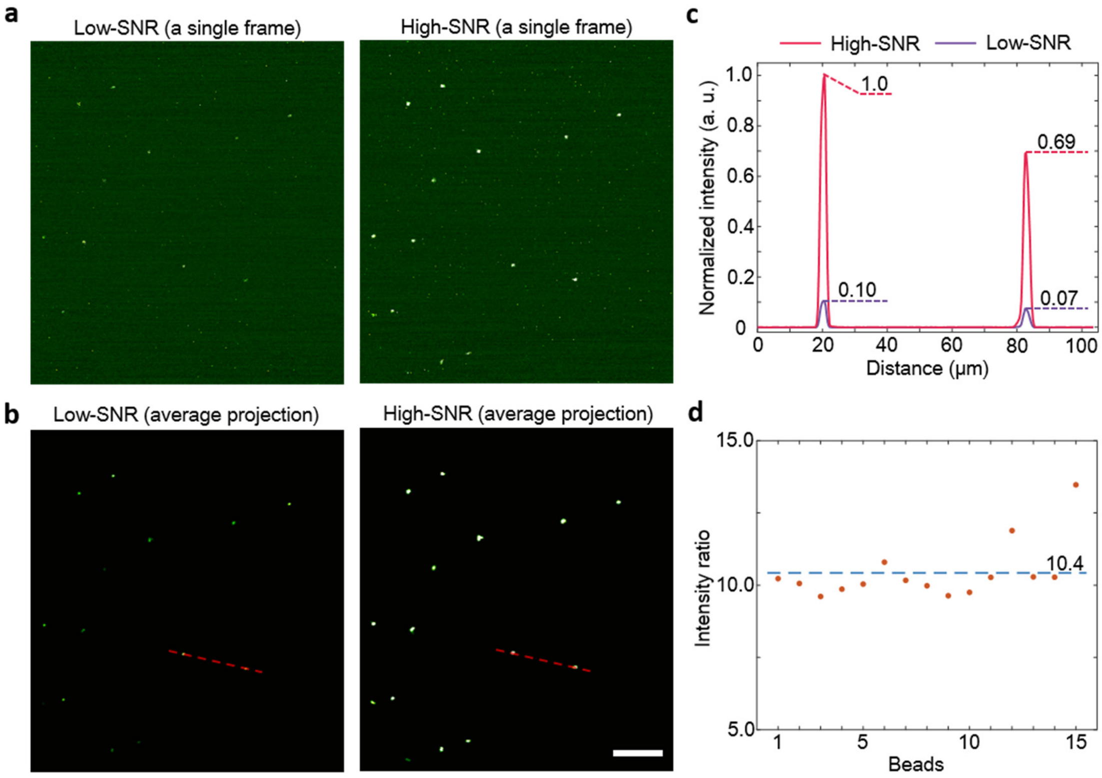


**Supplementary Figure 13**

**System calibration.**

**a**, Representative frames captured by the low-SNR detection path (left) and the high-SNR detection path (right). There were 15 isolated fluorescent beads (1 μm diameter) in the field of view (FOV). **b**, Average projection of 500 continuously acquired frames. Detection noise was largely suppressed and underlying fluorescence signals were revealed. **c**, Intensity profiles (normalized to the maximum of high-SNR recording) along the red dashed lines in **b**. Peak intensities are annotated. **d**, The intensity ratios (high-SNR relative to low-SNR) of all 15 fluorescent beads. Each point represents one bead. The average intensity ratio is 10.4 (blue dashed line).

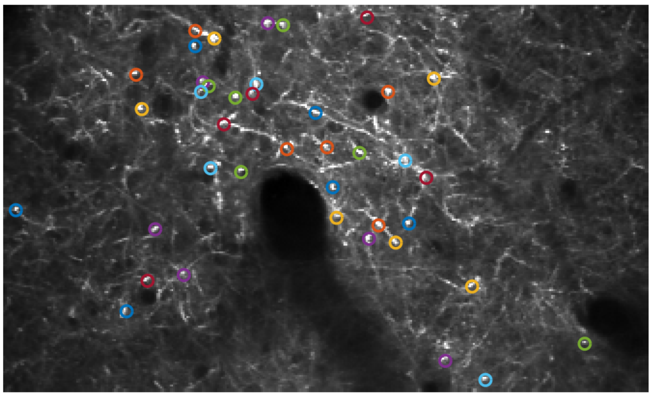

**Supplementary Figure 14**

**Pixels for extracting single-pixel fluorescence.**

Pixels (N=40) picked in neuropil imaging (Fig. 3a-c). Central pixels of the circles were used for extracting fluorescence traces.

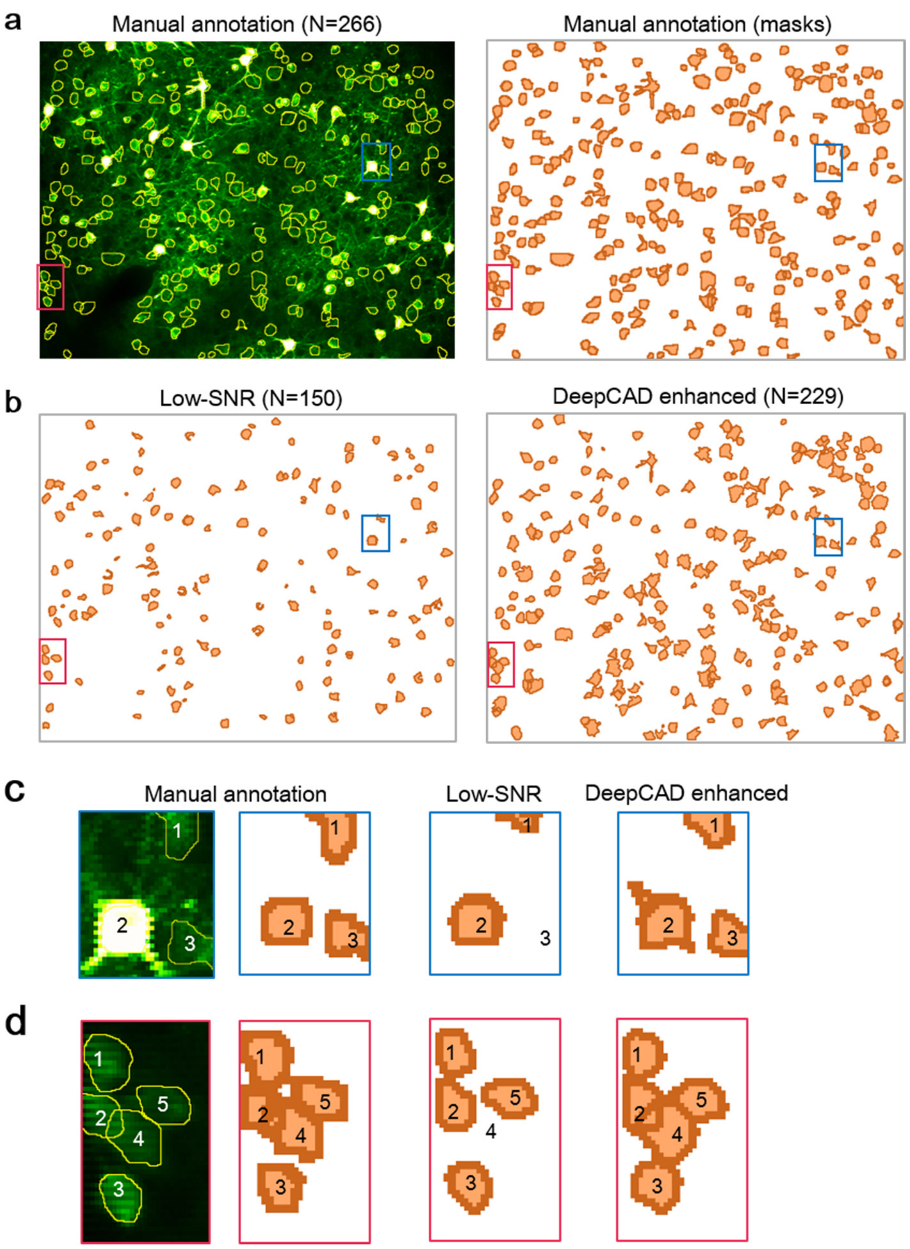


**Supplementary Figure 15**

**Human inspection of segmentation results.**

**a**, Left: manually annotated neuron borders. The standard deviation projection served as the background image. Right: manually annotated segmentation masks. **b**, Left: segmentation masks of the Low-SNR recording. Right: segmentation masks of the DeepCAD enhanced recording. Constrained nonnegative matrix factorization (CNMF) was used as the segmentation method. **c**, Magnified view of the blue boxed region shows the segmentation of three neurons. **d**, Magnified view of the red boxed region shows the segmentation results of five neurons. Missing neurons in the low-SNR recording can be recognized again after denoising.

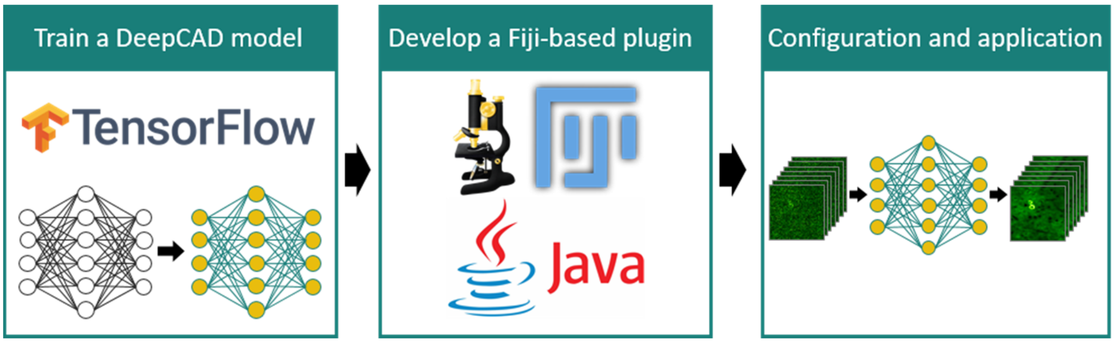


**Supplementary Figure 16**

**Fiji-based DeepCAD plugin.**

A Tensorflow implementation of DeepCAD was first trained using our self-supervised strategy. The pre-trained model was memorized in network parameters and saved as a separate file. Then, we created a Java project and packaged the application process of DeepCAD as a user-friendly plugin. Our plugin was based on the CSBDeep[11] framework of Fiji. Some necessary modifications were made for better performance and compatibility with our python code. The packaged DeepCAD model is out there to use after an easy configuration.

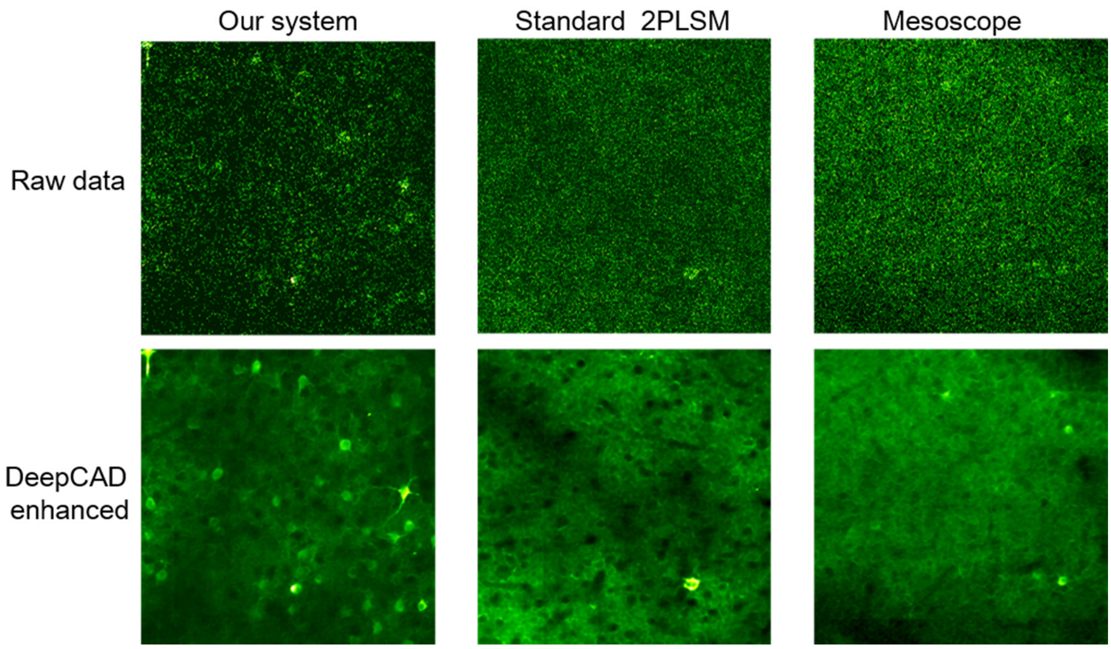


**Supplementary Figure 17**

**Cross-system validation.**

Denoising performance of DeepCAD on three two-photon laser-scanning microscopes (2PLSMs) with different system setups. Our system was equipped with alkali PMTs (PMT1001, Thorlabs) and a 25×/1.05 NA commercial objective (XLPLN25XWMP2, Olympus). The standard 2PLSM was equipped with a GaAsP PMT (H10770PA-40, Hamamatsu) and a 25×/1.05 NA commercial objective (XLPLN25XWMP2, Olympus). The two-photon mesoscope was equipped with a GaAsP PMT (H11706-40, Hamamatsu) and a 2.3×/0.6 NA custom objective. The same pre-trained model was used for denoising these data.

### Supplementary Notes

#### 1. Mixed Poisson-Gaussian Noise

Fluorescence microscopy is inherently sensitive to detection noise because the photon flux in fluorescence imaging is usually three orders of magnitude lower than that in photography[12]. Considering the detection physics, three sources mainly contribute to the detection noise in fluorescence imaging, namely the dark noise, the photon shot noise, and the readout noise[13]. Among them, the dark noise and the photon noise follow a Poisson distribution and the readout noise follows a Gaussian distribution. Hence, the detection noise of fluorescence microscopy follows a Mixed Poisson-Gaussian (MPG) distribution that can be formulated as

$$I_{acq} = P(I_{clean}) + N(0,\sigma^2), \tag{S1}$$

where $I_{clean}$ and $I_{acq}$ are the photon distribution (ground truth) and corresponding noise-contaminated photon distribution, respectively. The max values of them are scaled to 65535. $P$ is the function operator for the signal-related Poisson noise. $N$ is the signal-independent zero-mean Gaussian noise characterized by variance $\sigma^2$. Furtherly, digitization resolution (bit depth of the digitizer) is taken into consideration and the generation of MPG noise can be reformulated as

$$I_{acq} = MPG(I_{clean}; Q, n, \sigma^2) = \frac{2^n - 1}{Q} \cdot P\left(\frac{I_{clean} \cdot Q}{2^n - 1}\right) + N(0,\sigma^2). \tag{S2}$$

Here, $n$ is the bit depth ($n$ = 16 in our simulation) of captured images and is set to be 16 in most cases in microscopy. $MPG(\cdot)$ is the operator for MPG noise generation to simplify the mathematical expression. $Q$ is the relative photon number of each pixel captured by the detector. Different imaging signal-to-noise ratios (SNRs) are simulated with different $Q$.

Benefiting from advanced manufacturing technologies of integrated circuits and high-performance sensor cooling methods, the dark noise and the readout noise are relatively low. Thus, the photon shot noise plays the dominant role in the MPG noise of fluorescence microscopy. To approximate this property, we used a small $\sigma$ (1000 for

16-bit image) for the Gaussian component to keep the Poisson component dominant.

**2. Generation of simulated calcium imaging data**

Calcium imaging records calcium transients of large neural populations. Because of the invariance of the neuronal positions, fluorescence only fluctuates with neural activities. This process can be modeled as a matrix decomposition problem, which can be formulated as

$$S = A \cdot C, \tag{S3}$$

where $S$ is the time-lapse video composed of multiple frames, $A$ is the spatial components describing the positions of all neurons, $C$ is the temporal components describing the activities (without noise) of all neurons. If we specify the image dimension as $M \times N$, the number of frames as $T$, and the number of neurons recorded as $K$, then the dimensions of $S$, $A$, and $C$ are $(M \times N) \times T$, $(M \times N) \times K$, and $K \times T$, respectively. All 2D images are reshaped as 1D vectors. With the MPG noise defined in Eq.S2, the acquired noised-contaminated calcium imaging video can be generated using the following equation

$$\begin{aligned} S_{acq} &= MPG(S_{clean}; Q, n, \sigma^2) = MPG(A \cdot C; Q, n, \sigma^2) \\ &= \frac{2^n - 1}{Q} \cdot P\left(\frac{S_{clean} \cdot Q}{2^n - 1}\right) + N(0, \sigma^2) \end{aligned}, \tag{S4}$$

Here $S_{clean}$ and $S_{acq}$ are the clean video (ground truth) and corresponding noise-contaminated video, respectively. Because the noise of a certain pixel is only related to its intensity, the processes of video synthesis and noise addition can be conducted sequentially. The noise addition process can also be conducted frame by frame.

To synthesize realistic calcium imaging videos, we made a neuronal profile library containing spatial profiles of 517 neurons, which were extracted from a real two-photon imaging video with the constrained nonnegative matrix factorization[14,15] (CNMF) algorithm. Neurons were randomly selected from the library to form the spatial component $A$. For the temporal component $C$, we used MLspike[9], which can combine the physiological model of specified calcium indicators, to generate calcium traces for all neurons. The finally simulated data is synthesized using Eq. S4. The properties of simulated data are investigated in Supplementary Fig. 6.

#### 3. Tutorial of DeepCAD plugin

To ameliorate the difficulty of using our deep self-supervised learning-based method, we developed a user-friendly Fiji plugin (Supplementary Fig. 16). This plugin is easy to install and convenient to use. Researchers without expertise in computer science and machine learning can learn to use it in a very short time. The plugin is kept updating at the companion GitHub page. We provide a very short tutorial as follows.

##### 3.1 Install DeepCAD plugin on Fiji.

1. Download and install Fiji at: https://imagej.net/Downloads
2. Download the plugin file (DeepCAD.jar) from the companion GitHub repository (https://github.com/cabooster/DeepCAD).
3. Open Fiji and choose the plugin file (DeepCAD.jar) for installation (Plugins > Install…).
4. Restart Fiji and then you can found the DeepCAD plugin is installed and out there to use at: Plugins > DeepCAD.

##### 3.2 Use DeepCAD plugin for denoising of calcium imaging data.

1. Open Fiji.
2. Open the calcium imaging stack to be denoised.
3. Open the plugin at: Plugins > DeepCAD.
4. Select the pre-trained model and set six parameters on the panel (with default values and no changes are required unless necessary).
5. Click 'OK' and the denoised result will be displayed in another window after several minutes (depends on your data size).

##### 3.3 Train a customized DeepCAD model (optional).

Due to imaging systems and experiment conditions varies, a customized DeepCAD model trained on specified data is recommend for optimal performance. A Tensorflow implementation of DeepCAD is made publicly accessible at the companion GitHub repository of this paper (https://github.com/cabooster/DeepCAD) and a detailed document is provided to make it easy to use.
